# Patterning of the female reproductive tract along antero-posterior and dorso-ventral axes is dependent on *Amhr2+* mesenchyme in mice

**DOI:** 10.1101/2022.05.03.490514

**Authors:** Shuai Jia, Jillian Wilbourne, McKenna J Crossen, Fei Zhao

## Abstract

Morphogenesis of the female reproductive tract is regulated by the mesenchyme. However, the identity of the mesenchymal lineage that directs the patterning of the female reproductive tract has not been determined. Using in vivo genetic cell ablation, we identified *Amhr2*+ mesenchyme as an essential mesenchymal population in patterning the female reproductive tract. After partial ablation of *Amhr2*+ mesenchymal cells, the oviduct failed to develop its characteristic coiling due to decreased epithelial proliferation and tubule elongation during development. The uterus displayed a reduction in size and showed decreased cellular proliferation in both epithelial and mesenchymal compartments. More importantly, in the uterus, partial ablation of *Amhr2*+ mesenchyme caused abnormal lumen shape and altered the direction of its long axis from the dorsal-ventral axis to the left-right axis (i.e. perpendicular to the dorsal-ventral axis). Despite these morphological defects, epithelia underwent normal differentiation into secretory and ciliated cells in the oviduct and glandular epithelial cells in the uterus. These results demonstrated that *Amhr2*+ mesenchyme can direct female reproductive tract morphogenesis by regulating epithelial proliferation and lumen shape without affecting the differentiation of epithelial cell types.

## Introduction

Establishment of the female reproductive tract organs requires proper morphogenesis of the Müllerian duct, the progenitor for the epithelium of the entire female reproductive tract organs. Along the antero-posterior (i.e. rostral-caudal) axis, the Müllerian duct gives rise to structurally and functionally unique organs including the coiled oviduct, the straight uterus, cervix and the upper vagina (Kurita, 2011; Mullen and Behringer, 2014). Classic tissue recombination studies have demonstrated that organ-specific mesenchymal signaling dictates the regionalization of the female reproductive tract along the antero-posterior axis. For instance, tissue recombinants consisting of uterine mesenchyme and vaginal epithelium displayed uterine morphology; while the tissue recombinants consisting of vaginal mesenchyme and uterine epithelium underwent vaginal morphogenesis (Cunha, 1976). The female reproductive tract, particularly the uterus, is also patterned along the dorso-ventral axis (also known as the mesometrial-anti-mesometrial axis) (Dey et al., 2004). The uterine epithelial lumen is elongated along the dorsal-ventral axis at birth and forms differential structures at the dorsal and ventral sides. Epithelia at the dorsal (mesometrial) side develops the uterine rail, a morphological structure similar to a railroad track along the dorsal pole (Vue et al., 2018). On the other hand, at the ventral (anti-mesometrial) side, luminal epithelium forms the ventral ridge, a continuous and smooth longitudinal epithelial invagination into the surrounding mesenchyme to establish uterine glands (Vue and Behringer, 2020). As a result, uterine gland formation is absent on the dorsal side and only occur at the ventral region (Kelleher et al., 2019). The differential establishment of uterine glands along the dorso-ventral axis is regulated by differential mesenchymal WNT signaling. Active Wnt signaling that is essential for uterine gland formation is inhibited by the Wnt inhibitor *Dkk2* derived specifically from the dorsal mesenchyme (Goad et al., 2017). These observations show that patterning of female reproductive tract organs along both the antero-posterior and dorso-ventral axes is governed by the mesenchyme.

AMHR2 (Anti-Müllerian hormone receptor type II) is the specific receptor for mediating the action of anti-Müllerian hormone (AMH), a member of the transforming growth factor beta gene family (Mullen and Behringer, 2014). *Amhr2* is specifically expressed in Müllerian duct mesenchyme in both male and female embryos around E12.5-E13 (Arango et al., 2008; Kobayashi et al., 2011). In male embryos, AMHR2 is activated by testis-derived AMH to induce Müllerian duct regression via mesenchymal-epithelial interaction (Arango et al., 2008; Kobayashi et al., 2011). On the other hand, in female embryos, the ovary does not produce AMH; as a result, AMHR2 mediated pathway remains inactive, and the Müllerian duct is maintained. Despite the lack of the ligand activation, *Amhr2* expression is still present and displays dynamic pattern in the mesenchyme in females. *Amhr2* is predominantly expressed in the prospective ventral (anti-mesometrial) mesenchyme at the fetal stage and becomes restricted to a thin layer of subluminal mesenchyme in the uterus after birth before PND6. When the myometrium is formed from PND6 onwards, its expression is detected only in the circular smooth muscle layer (Arango et al., 2008; Saatcioglu et al., 2019). When *Amhr2* promoter driven Cre (*Amhr2-Cre*) was generated and crossed with a reporter, the majority of mesenchymal cells in the oviduct and uterus were labeled starting from E12.5 (Huang et al., 2012; Kobayashi et al., 2011). Therefore, *Amhr2-Cre* has been widely used to study functions of the mesenchymal factors in female reproductive tract biology. As reported in previous literature, deletion of either *Tgfbr1* or *Tbfbr2* in *Amhr2+* mesenchyme caused oviductal diverticula, myometrial defects, and endometrial hyperplasia (Li et al., 2011; Ni et al., 2021). Ablations of the key Hippo kinases *Lats1* and *Lats2* in the *Amhr2*+ mesenchyme led to profound developmental defects including cystic oviducts, shortened uterine horns, and the absence of uterine glands (St-Jean et al., 2019).

Targeting a couple of genes at a time does not reveal the comprehensive role of *Amhr2*+ mesenchyme in female reproductive tract development. A powerful way to elucidate the function of a specific cell population within the whole organism is in vivo genetic cell ablation using the *Rosa-DTA* (diphtheria toxin fragment A) mouse line. Upon Cre-mediated excision of the *loxP*-flanked transcriptional stop sequence, DTA expression is induced specifically in Cre-expressed cells, where DTA induces cell apoptosis by inhibiting protein translation (Ivanova et al., 2005). Because mice do not express the membrane receptor for the internalization of DTA, DTA released from the apoptotic cells cannot enter surrounding non-Cre-expressed cells, avoiding non-specific cell ablation (Ivanova et al., 2005). We therefore set out to use *Amhr2-Cre; Rosa-DTA* genetic cell ablation model to specifically ablate *Amhr2+* mesenchyme to reveal functional significances of *Amhr2+* mesenchyme in female reproductive tract development.

## Materials and Methods

### Mice

*Amhr2-Cre* line was provided by Richard Behringer (The University of Texas MD Anderson Cancer Center) and maintained in C57BL/6 genetic background (Jamin et al., 2002). *Rosa-DTA* line (stock# 006331) was originally purchased from Jackson Laboratories (Bar Harbor, ME) and maintained with their original genetic background (C57BL/6J and CD1). Timed mating was set up by housing stud males with 2-3 sexually mature females (2 to 6 months old) in the later afternoon. Vaginal plugs were checked early on the next morning, and the day of plug detection was designated as embryonic day 0.5 (E0.5). The day of birth was determined by examining pup delivery in the afternoon and was designated as postnatal day 0 (PND0). The embryos or mice harboring both *Amhr2^Cre/+^* and *Rosa^DTA/+^* were used in the ablation group while mice of other genotypes (*Amhr2^Cre/+^;Rosa^+/+^, Amhr2^+/+^;Rosa^DTA/+^,* or *Amhr2^+/+^;Rosa^+/+^*) were used as the control. All experiments were performed at least on three mice for each group and the number of mice used for each experiment was described in figure legends. All mouse procedures were approved by the University of Wisconsin-Madison (UW-Madison) Animal Care and Use Committees and are in compliance with UW-Madison approved animal study proposals and public laws.

### Genotyping

PCR was performed with primers (Forward: 5’-CGCATTGTCTGAGTAGGTGT-3’, Reverse: 5’-GAAACGCAGCTCGGCCAGC-3’) for *Amhr2-Cre* allele (Li et al., 2008) and primers (Mutant Reverse: 5’-GCGAAGAGTTTGTCCTCAACC-3’, Common: 5’-AAAGTCGCTCTGAGTTGTTAT-3’, Reverse: 5’-GGAGCGGGAGAAATGGATATG-3’) for *Rosa-DTA* allele. Platinum™ II Taq Hot-Start DNA Polymerase (Invitrogen™, Catalog #: 14966001) was used to run the thermal cycle: 94°C for 2 min, 34 cycles of [94°C for 15 sec, 60°C for 5 sec, and 68°C for 15 sec] followed by 68°C for 5 min.

### Tissue processing, embedding and sectioning

Tissues were fixed in 10% neutral buffered formalin (Leica, Catalog #: 3800598) overnight at room temperature, washed 3 times with 1xPBS for 10 min each, dehydrated by a serial of ethanol (70%, 80% and 95% for 30 min for each; 100% ethanol I and II for 50 min for each), cleared (100% ethanol: xylene =1:1 for 5 min; xylene I for 5 min; xylene II for 3 min) and infiltrated with paraffin (100% ethanol: soft paraffin =1:1 for 0.5 h; soft paraffin I for 1.5 h; soft paraffin II for 1 h; soft paraffin: hard paraffin=1:1 for 1 h; hard paraffin for 1.5 h) before being embedded in paraffin. Tissues were sectioned at 5 μm using a microtome (TN6000, Tanner Scientific) and were collected at least 50 µm apart from each other for subsequent immunofluorescence staining and quantifications.

**H&E staining** was performed as previously described (Zhao et al., 2013). Briefly, paraffin sections were deparaffinized, rehydrated and stained with hematoxylin (Electron Microscopy Sciences, Catalog #: 26123-08). After hematoxylin staining, slides were rinsed in running tap water, decolorized in acid ethanol, rinsed in running tap water and then counterstained with eosin (Leica Biosystems, Catalog #: 3801615). After counterstaining, sections were dehydrated with gradient ethanol, cleared in xylene and mounted with mounting medium (Fisher Chemical, Catalog #: SP15500). All images were taken under a Nikon ECLIPSE E600 microscope.

### Immunofluorescence

Paraffin sections were deparaffinized and rehydrated (the same as described in H&E staining). Sections were then subjected for antigen retrieval using a microwave oven (35PF91, Grainger) with commercial antigen unmasking solution (VECTOR, Catalog #: H-3300) and underwent immunostaining procedure using the Sequenza™ Immunostaining Center System (Electron Microscopy Sciences, Catalog #: 71407-01)(Zhao et al., 2017). Briefly, sections were washed with 1xPBS and PBST (1x PBS with 0.1% Triton X-100) and incubated with blocking solution (5% Donkey serum in PBST) for 1 h at room temperature and then with primary antibody in blocking solution overnight at 4°C. After being washed three times in PBST, sections were incubated with secondary antibodies for 1 h at room temperature and counterstained by DAPI (1:1000, Thermo Scientific™, Catalog #:62248), coverslipped with EverBrite mounting medium (Biotium, Catalog #: 23003) for imaging under a Leica TCS SP8 confocal microscope.

The following primary antibodies were used: rabbit anti-Cleaved PARP1 [E51] (1:300, Abcam, Catalog #: ab32064), rabbit anti-Ki67(1:300, Abcam, Catalog #: ab15580), rabbit anti-alpha-smooth muscle actin (αSMA) (1:200, Abcam, Catalog #: ab5694), rabbit anti-FOXA2 [EPR4466] (1:200, Abcam, Catalog #: ab108422), rabbit anti-PAX8 polyclonal (1:200, Proteintech, Catalog #: 10336-1-AP), and mouse anti-β-Tubulin IV [Clone ONS1A6] (1:200, Biogenex, Catalog #: MU178). The secondary antibodies conjugated with different fluorescent dyes were used: donkey anti-rabbit IgG (H+L), Alexa Fluor 488 (1:200, Invitrogen, Catalog #: A-21206) and donkey anti-mouse IgG (H+L), Alexa Fluor 568 (1:200, Invitrogen, Catalog #: A-10037).

### Quantifications of cleaved-PARP1+ and Ki67+ cells

Images of 3-7 sections from each of the 5-7 mice were analyzed using ImageJ. For quantifying cleaved-PARP1+ cells, autofluorescent cells identified by imaging in another color channel were excluded (Fayzullina and Martin, 2014). For Ki67+ proliferating cells, only cells with over 50% of the nucleus exhibiting Ki67 positive staining were counted as cells in the active process of mitosis (Miller et al., 2018; Saadeh et al., 2020). To count Ki67+ cells in the mesenchymal compartment, images of uterine sections were divided into dorsal and ventral regions by drawing lines perpendicular to the dorso-ventral axis at the center of epithelial lumen. The total number of cells (DAPI staining for individual nucleus), cleaved-PARP1+ and Ki67+ cells in epithelial or mesenchymal compartments in each section were counted twice and averaged. The percentage of apoptotic or proliferating cells was calculated as the fraction of cleaved-PARP1+ or Ki67+ cells over the total number of cells in each section for subsequent statistical analysis. Three independent examiners followed the same criteria and procedures for counting cells to confirm statistical differences.

### Quantifications of FOXA2+ uterine glands

Images of 3-5 sections from PND21 uteri were analyzed using ImageJ.

### Measurement of oviductal length

After removing connective tissues, the oviduct was placed on a membrane for image acquisition. In the image, a curve was artificially drawn at the center of the oviductal lumen from the distal end (the ostium) of infundibulum to the uterine-tubal junction (the flexura medialis on E16.5) (Agduhr, 1927). ImageJ was used to measure the length of the curve twice and average the two measurements for each oviduct. Three independent examiners followed the same criteria and procedures to confirm statistical differences.

### Statistical analyses

The significant differences between control and ablation groups were evaluated by two-tailed unpaired Student’s t-test using GraphPad Prism 9.0 software. Results were shown as mean ± s.e.m. The significance level was set at p<0.05.

## Results

### The number of apoptotic cells in the mesenchymal compartment was significantly increased in the *Amhr2-Cre; Rosa-DTA* female reproductive tract

The Cre activity of *Amhr2-Cre* initiates around E12.5 in the Müllerian duct mesenchyme (Kobayashi et al., 2011). It takes up to 48h from the onset of Cre expression to the recombination of the *loxP*-flanked stop codon, allowing DTA expression and DTA-induced apoptosis (Brockschnieder et al., 2004; Ivanova et al., 2005). In addition, the antero-posterior regionalization of the Müllerian duct occurs around E16.5 (Huang et al., 2014; Ma et al., 1998; Stewart and Behringer, 2012a). We therefore examined the expression of apoptotic marker, cleaved Poly [ADP-ribose] polymerase 1 (PARP1) in E16.5 Müllerian ducts to determine whether our *Amhr2-Cre; Rosa-DTA* model successfully induced death of *Amhr2+* mesenchymal cells (D’Amours et al., 2001) (**Fig. 1A-1H**). By correcting background autofluorescent cells, we found that less than 0.1% cells in mesenchymal compartment were positive for cleaved-PARP1 in the anterior (future oviduct, 0.02%±0.02%) and posterior (future uterus, 0.07%±0.06%) Müllerian ducts of the control group. On the contrary, percentages of cleaved-PARP1+ cells were significantly increased both in the anterior (0.34%±0.05%) and posterior regions (0.95%±0.12%) of the ablation group (**Fig. 1I**). When *Amhr2-Cre* reporter lines were used to track *Amhr2*+ mesenchyme (Kobayashi et al., 2011; Zhao et al., 2022), *Amhr2*+ mesenchymal cells contributed to the majority of mesenchymal cells in both oviducts and uteri. The average percentage of apoptotic cells was less 1% in the mesenchymal compartment in the ablation group (**Fig. 1I**). These results demonstrate that the *Amhr2-Cre; Rosa-DTA* model successfully achieves partial ablation of *Amhr2+* mesenchymal cells.

**Fig 1.**
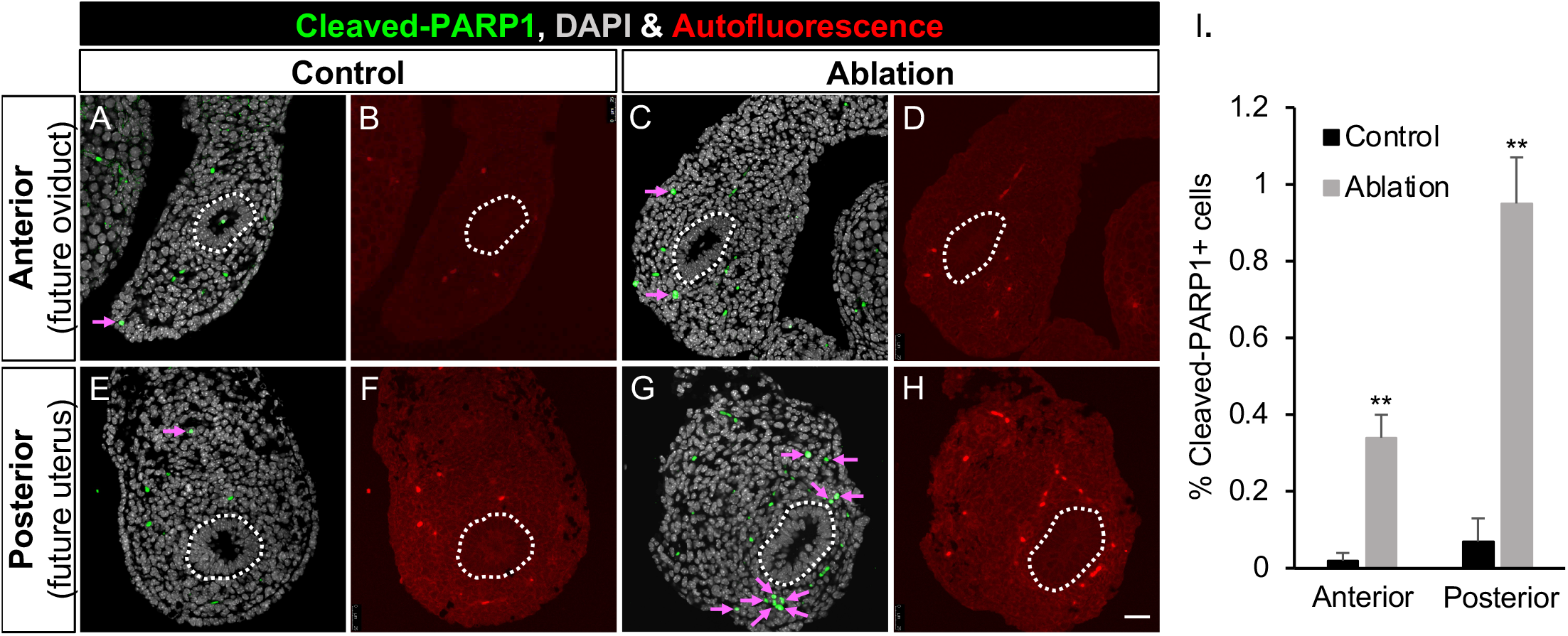
Increased number of apoptotic cells in the mesenchymal compartment of E16.5 Müllerian ducts in the ablation group. (A, E, C, G) Immunofluorescence staining of the apoptotic marker cleaved-PARP1 and DAPI for nuclei in the anterior and posterior regions of E16.5 Müllerian duct in the control (A & E) and ablation (C & G) groups. Pink arrows indicate identified apoptotic cells. (B, D, F, H) Autofluorescent cells identified by another color channel in the anterior and posterior regions of E16.5 Müllerian duct in the control (A & E) and ablation (C & G) groups. Dashed lines: Müllerian ducts. (I) Percentages of apoptotic cells in the control and ablation groups. Results were shown as mean ± s.e.m. and analyzed by two-tailed unpaired Student’s t test. **, p<0.01 compared to the control group at the same age. N=3 for each group. Scale bar: 25 µm.

### Defective oviductal coiling but normal oviductal epithelial regionalization and differentiation in the ablation group

Next, we determined the consequence of partial ablation of *Amhr2*+ mesenchymal cells by examining gross morphology of the female reproductive tract on PND21, when the patterning of the female reproductive tract is complete (Cooke et al., 2013). The female reproductive tracts were smaller and shorter in the ablation group compared to those in the control group (**Fig. 2A**). In the control group, the Müllerian duct differentiates into two morphologically distinct organs: the highly-coiled oviduct in the anterior region and the straight uterus in the posterior region (**Fig. 2A**). However, we found that the antero-posterior patterning was defective in the ablation group where the oviduct lost its characteristic coiling (**Fig. 2A**). In addition to coiling formation, the oviduct undergoes regionalization and further differentiation into three morphologically and histologically distinct segments along the antero-posterior axis: the infundibulum that has the ostium (the opening) surrounded fimbriae (fingerlike epithelial projections); the ampulla that possess more longitudinal epithelial folds but thin smooth muscle layers; and the isthmus that has less epithelial folds but thicker smooth muscle layers (Ruberte et al., 2017) (**Fig. 2B**). These three segments with their typical histological structures were observed in the uncoiled oviduct of the ablation group in both whole mount images and histological sections (**Fig. 2A & 2B**). The observation of uncoiled oviducts with normal regionalization suggests that oviductal coiling and regionalization could be two uncoupled events.

**Fig 2.**
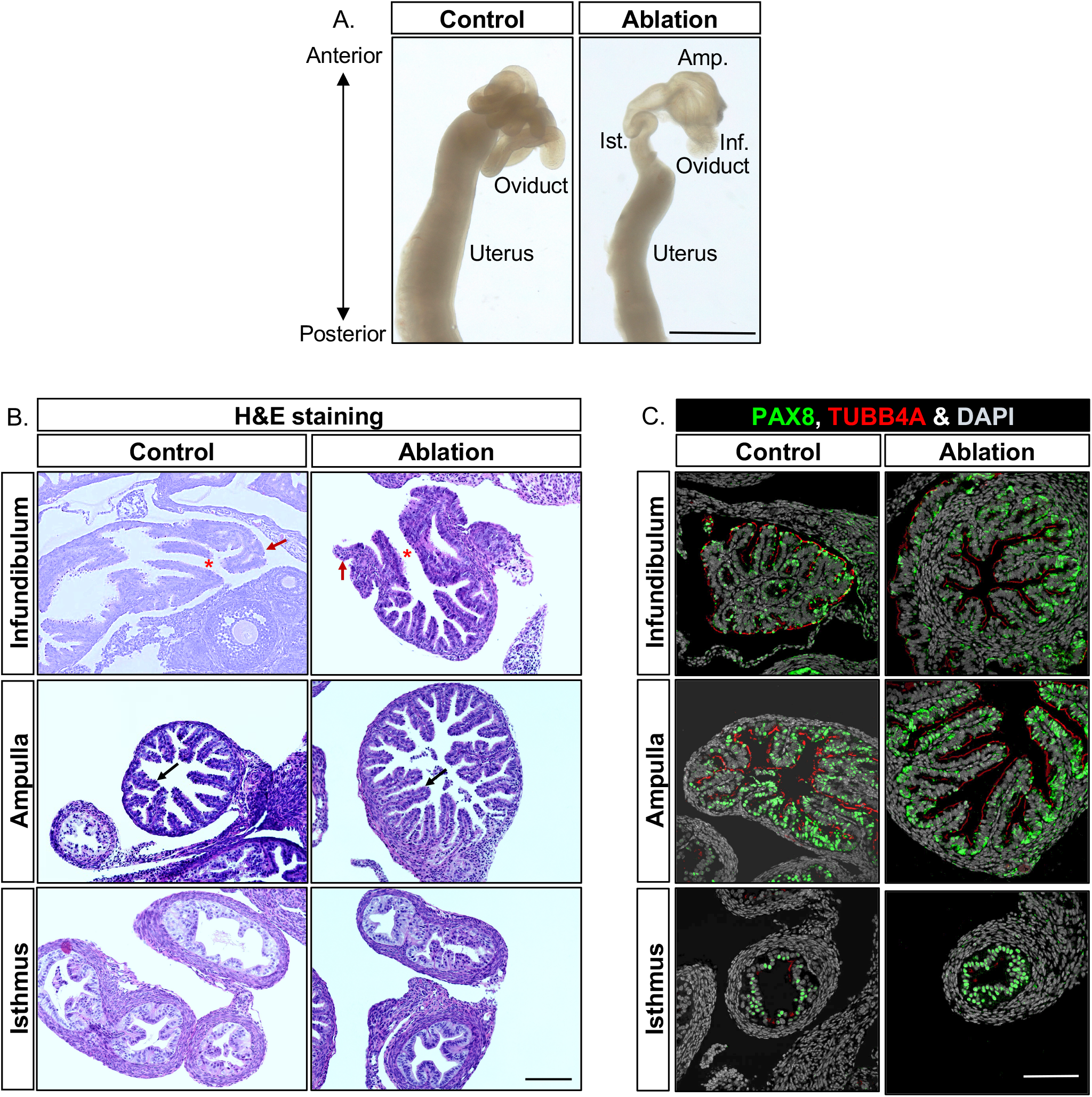
Defective oviductal coiling but normal epithelial regionalization and differentiation in the ablation group. (A) Representative images of PND21 oviducts and uteri of the control and ablation group. Scale bar: 2 mm. (B) Histology of PND21 oviducts and uteri of the control and ablation groups. Stars: ostium. Red and black arrows: fimbriae and epithelial folds, respectively. (C) Double immunofluorescence staining of the ciliated cell marker TUBB4A and secretory cell marker PAX8 in the infundibulum, ampulla, and isthmus. Scale bar: 100 µm in B & C. N=3 for both the control and ablation groups for A, B, C.

Oviductal epithelium consists of two major cell types, ciliated and secretory epithelial cells (Barton et al., 2020). To determine whether epithelial cell differentiation is impaired in the oviduct of the ablation group, we performed double-immunofluorescence staining of PAX8 (Paired box 8, a secretory epithelial cell marker) and TUBB4A (Tubulin beta 4A class IVa, a ciliated cell marker) to visualize both cell types. In the control group, PAX8 expression was detected in all three regions and its abundance was gradually increased from the infundibulum to the isthmus (**Fig. 2C**). On the other hand, TUBB4A+ ciliated cells were the most abundant in the epithelium of the infundibulum; and were decreased from the infundibulum to the isthmus (**Fig. 2C**). These results were consistent with other studies on the abundancy of ciliated and secretory cells from the infundibulum to the isthmus (Ford et al., 2021; Ghosh et al., 2017; Yuan et al., 2021). In the ablation group, both PAX8 and TUBB4A expression were detected in all three regions (**Fig. 2C**) and their abundances along the antero-posterior axis were comparable with those in the control group. These results demonstrated that the partial ablation of *Amhr2*+ mesenchymal cells leads to defective oviductal coiling without disrupting the normal epithelial regionalization and differentiation.

### Defective ovarian development in the ablation group

*Amhr2-Cre* also targets somatic cells in the ovary as early as E17.5 (Jorgez et al., 2004). We therefore determined whether ovarian development was affected on PND21. In the control group, ovaries were in oval shape and displayed follicles on the surface. In contrast, ovaries in the ablation group were irregular in shape and much smaller in size with few follicles on the surface (**Fig. S1**). These results demonstrate that ovarian and follicular development were impaired in our ablation model. However, it has been well-established that ovaries do not play any significant roles in fetal and postnatal female reproductive tract development before PND25 (Couse and Korach, 1999; Luo et al., 1994; Ogasawara et al., 1983). Therefore, abnormal ovarian development would not contribute to the phenotype of the female reproductive tract in our cell ablation model. We next focused on investigating cellular mechanisms of defective oviductal coiling observed in the ablation group.

### Impaired oviductal elongation during development in the ablation group

Coiling morphogenesis depends on the elongation of tubular organs (Savin et al., 2011). To determine whether oviductal elongation was affected during development, we measured the oviductal distance from the distal end of the oviduct to the uterine-tubal junction over the course of the oviducal coiling (E16.5-PND15) (Stewart and Behringer, 2012a). In the control group, the oviduct was a straight tube on E16.5, and then became coiled with an increased number of turns and lengths starting from PND0 (E16.5: 9.02 ± 0.28 mm; PND0: 11.99 ± 0.20 mm; PND3: 14.48± 0.97 mm; PND7: 22.27± 0.79 mm; PND15: 43.80± 1.49 mm) (**Fig. 3A & 3B**). On the contrary, in the ablation group, the oviduct remained uncoiled on all examined timepoints (**Fig. 3A**). On E16.5 prior to the initiation of coiling on PND0, the length of oviductal region in the ablation group was already significantly shorter than those of the control group. From E16.5 to PND15, the oviductal length in the ablation group was increased but remained 40-60% of their counterparts in the control group (E16.5: 5.54 ± 0.24 mm vs 9.02 ± 0.28 mm; PND0: 5.68± 0.19 mm vs 11.99 ± 0.20 mm; PND3: 8.38 ± 0.53 mm vs 14.48 ± 0.97 mm; PND7: 9.52 ± 1.30 mm vs 22.27 ± 0.79 mm; PND15: 21.41 ± 0.66 mm vs 43.80 ± 1.49 mm) (**Fig. 3B**). These results demonstrate that the partial ablation of *Amhr2*+ mesenchymal cells impairs oviductal elongation during development, which contribute to defective oviductal coiling.

**Fig 3.**
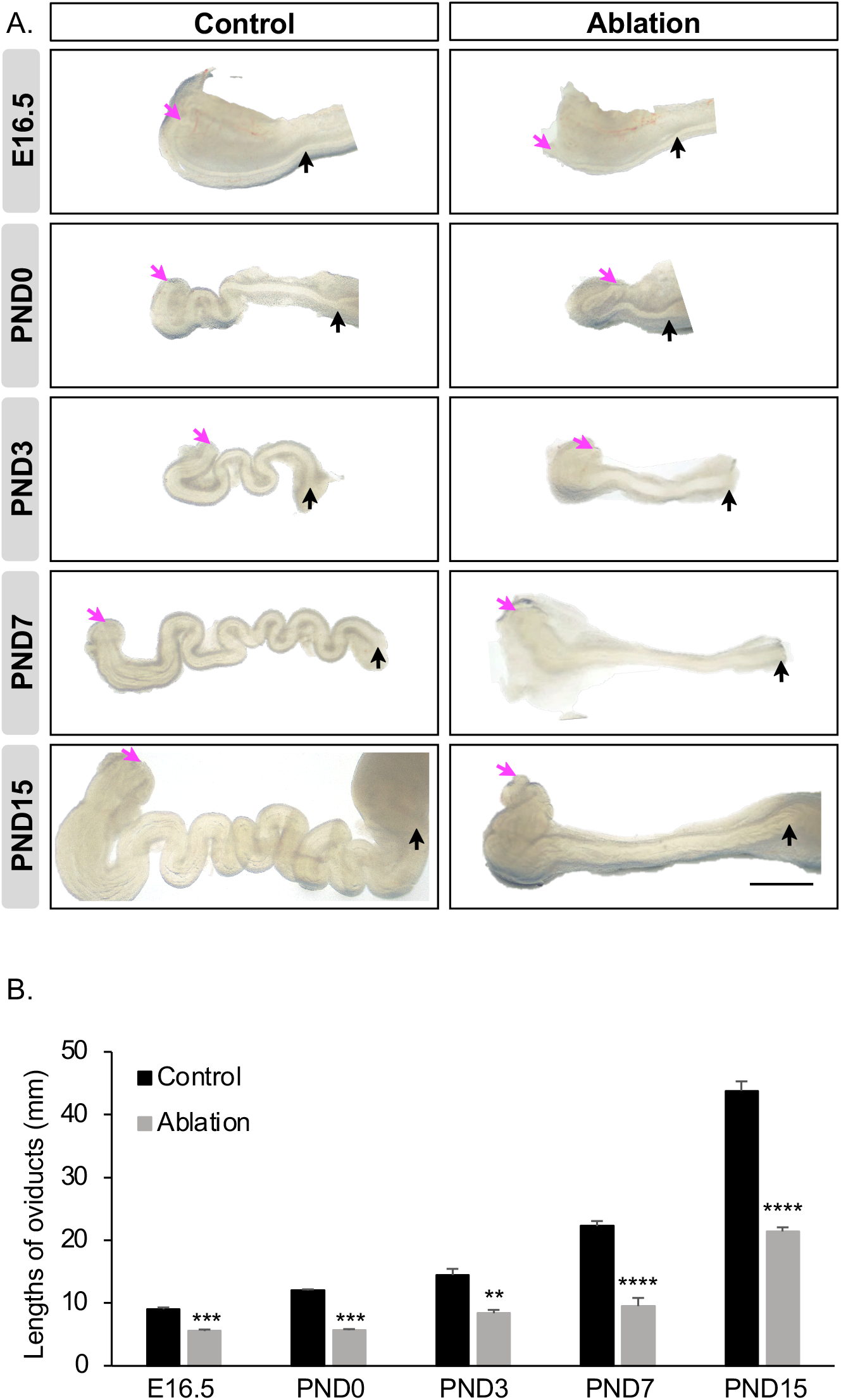
Impaired oviductal elongation during development in the ablation group. (A) Representative images of oviducts in the control and ablation groups from E16.5 to PND15. Scale bar: 1 mm. Pink and black arrows indicate the ostium of infundibulum and the uterine-tubal junction (the flexura medialis on E16.5), respectively. (B) Quantification of oviductal lengths in the control and ablation groups. N=3-7. Results were shown as mean ± s.e.m. and analyzed by two-tailed unpaired Student’s t test. **, p<0.01; ***, p <0.001; ****, p <0.0001 compared to the control group at the same age.

### Decreased proliferation in the oviductal epithelium in the ablation group

Epithelial proliferation has been implicated in tubular organ elongation (Andrew and Ewald, 2010). Therefore, we performed immunofluorescence staining of the proliferating marker Ki67 (Marker of Proliferation Ki-67) to determine the percentage of Ki67+ proliferating cells in oviductal epithelia on PND0 when the oviductal elongation initiates (**Fig. 4A**). In the control group, the percentage of Ki67+ cells was 30% on the average (**Fig. 4B**), which was significantly decreased to 20% in the ablation group. These results demonstrate that the partial ablation of *Amhr2*+ mesenchymal cells significantly decreases oviductal epithelial proliferation.

**Fig 4.**
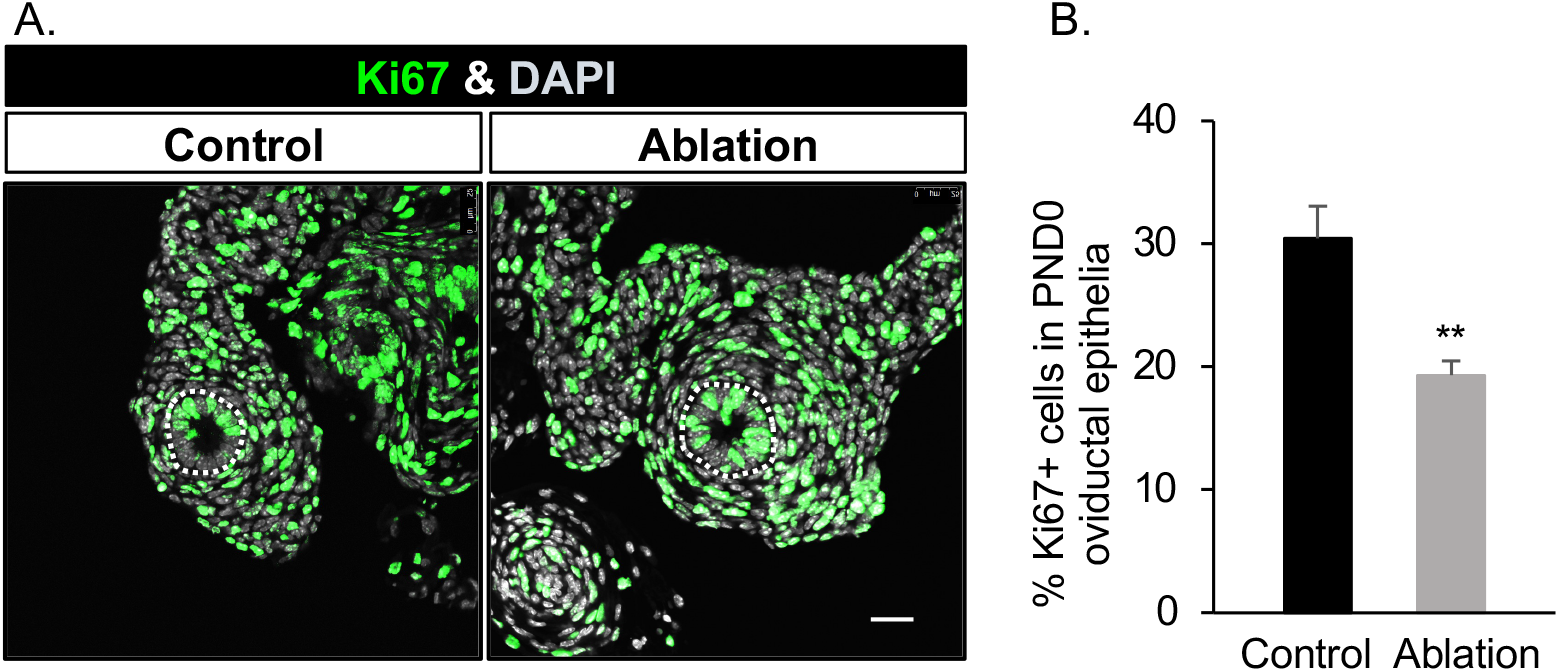
Decreased proliferation in the oviductal epithelium in the ablation group. (A) Representative images of Ki67 immunofluorescence staining on PND0 oviducts in the control and ablation groups. White dashed circles: oviductal epithelium. Scale bar: 25 µm. (B) Percentages of Ki67+ cells in the PND0 oviductal epithelia of the control and ablation groups. N=6, 5 for the control and ablation group, respectively. Results were shown as mean ± s.e.m. and analyzed by two-tailed unpaired Student’s t test. **, p<0.01.

### Normal smooth muscle formation in the oviduct of the ablation group

Smooth muscles provide mechanical forces in tubular organ morphogenesis (Jaslove and Nelson, 2018). We next investigated whether smooth muscle formation in the oviduct was affected in the ablation group by performing immunofluorescence staining of the smooth muscle marker αSMA on PND3, when smooth muscle cells initiate their differentiation, and PND21 (Agduhr, 1927; Stewart and Behringer, 2012b) (**Fig. S2**). In both the control and ablation groups, αSMA expression was detected in the inner circular layer of mesenchymal compartments in both anterior and posterior oviductal sections on PND3 (**Fig. S2A)** and PND21 **(Fig. S2B**). In addition, αSMA stained mesenchymal layer was thicker in the isthmus region than that in the ampulla regions as seen in the normal condition (**Fig. S2B**). These results demonstrate that partial ablation of *Amhr2*+ mesenchymal cells does not affect smooth muscle formation in the oviduct.

### Defective uterine patterning along the dorsal-ventral axis in the ablation group

To determine the consequence of partial ablation of *Amhr2*+ mesenchymal cells on uterine development, we examined uterine sections on PND0 and PND21, during which the epithelium undergo extensive morphogenesis and forms uterine glands (Brody and Cunha, 1989). In the control group, uterine lumen was elongated along the dorsal-ventral axis on PND0, forming a flattened circle in the cross-section view (**Fig. 5A**). The long axis of the uterine lumen remained along the dorsal-ventral axis on PND21 (**Fig. 5C**). However, in the ablation group, the size of uterine cross-sections was much smaller. More importantly, lumen shape became irregular with its long axis perpendicular to the dorsal-ventral axis on both PND0 and PND21(**Fig. 5B & 5D**).

**Fig 5.**
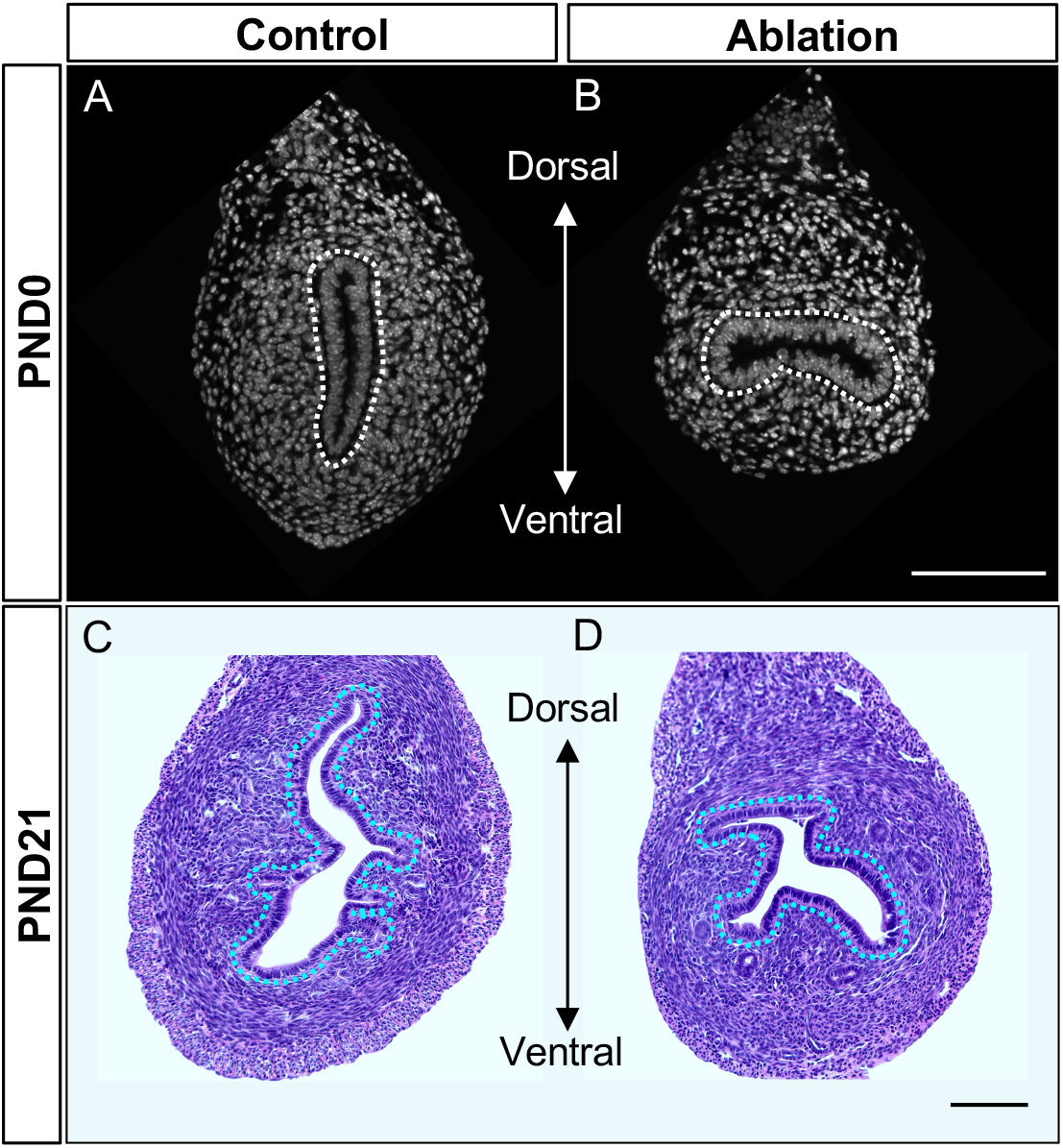
Defective uterine patterning along the dorsal-ventral axis in the ablation group. (A & B) Representative images of DAPI staining of PND0 uteri in the control and ablation groups. (C & D) Histology of PND21 uteri in the control and ablation groups. Dashed circles: uterine lumens. Scale bar: 100 µm. N=6, 5, 4, 3 for A, B, C, D, respectively.

On PND21, the uterine epithelium extended into the stroma and established uterine glands at the lateral and ventral regions (**Fig. S3**). In the ablation group, uterine glands were also observed in these two regions, and the number of uterine glands was decreased without significant differences (**Fig. S3**). These results indicate that uterine gland formation and patterning was not affected in the ablation group. Taken together, these results demonstrate that partial ablation of *Amhr2*+ mesenchymal cells does not affect uterine gland formation but disrupts the patterning of uterine epithelial lumen along the dorsal-ventral axis.

### Decreased proliferation in epithelial and mesenchymal compartments of the uterus in the ablation group

We then performed immunostaining of the proliferating marker Ki67 to determine the percentage of Ki67+ proliferating cells in uterine epithelium and mesenchyme on PND0 when uterine defects were observed (**Fig. 6**). In the control group, average percentages of Ki67+ in epithelia and the whole mesenchyme were 26% and 27%, respectively (**Fig. 6B**). However, the percentages were significantly decreased to 16% and 20% in the ablation group (**Fig. 6**). Given patterning of uterine lumen along the dorsal-ventral axis was altered, we then determined whether percentages of Ki67+ proliferating cells were differential in uterine dorsal and ventral mesenchyme. We found that percentages of Ki67+ proliferating cells in both dorsal and ventral regions were decreased but with statistical significance only observed in the dorsal mesenchymal compartments (**Fig. 6B**, P=0.02 on the dorsal side; P=0.07 on the ventral side). These results demonstrate that partial ablation of *Amhr2*+ mesenchymal cells decreases cellular proliferation in epithelial and mesenchymal compartments.

**Fig 6.**
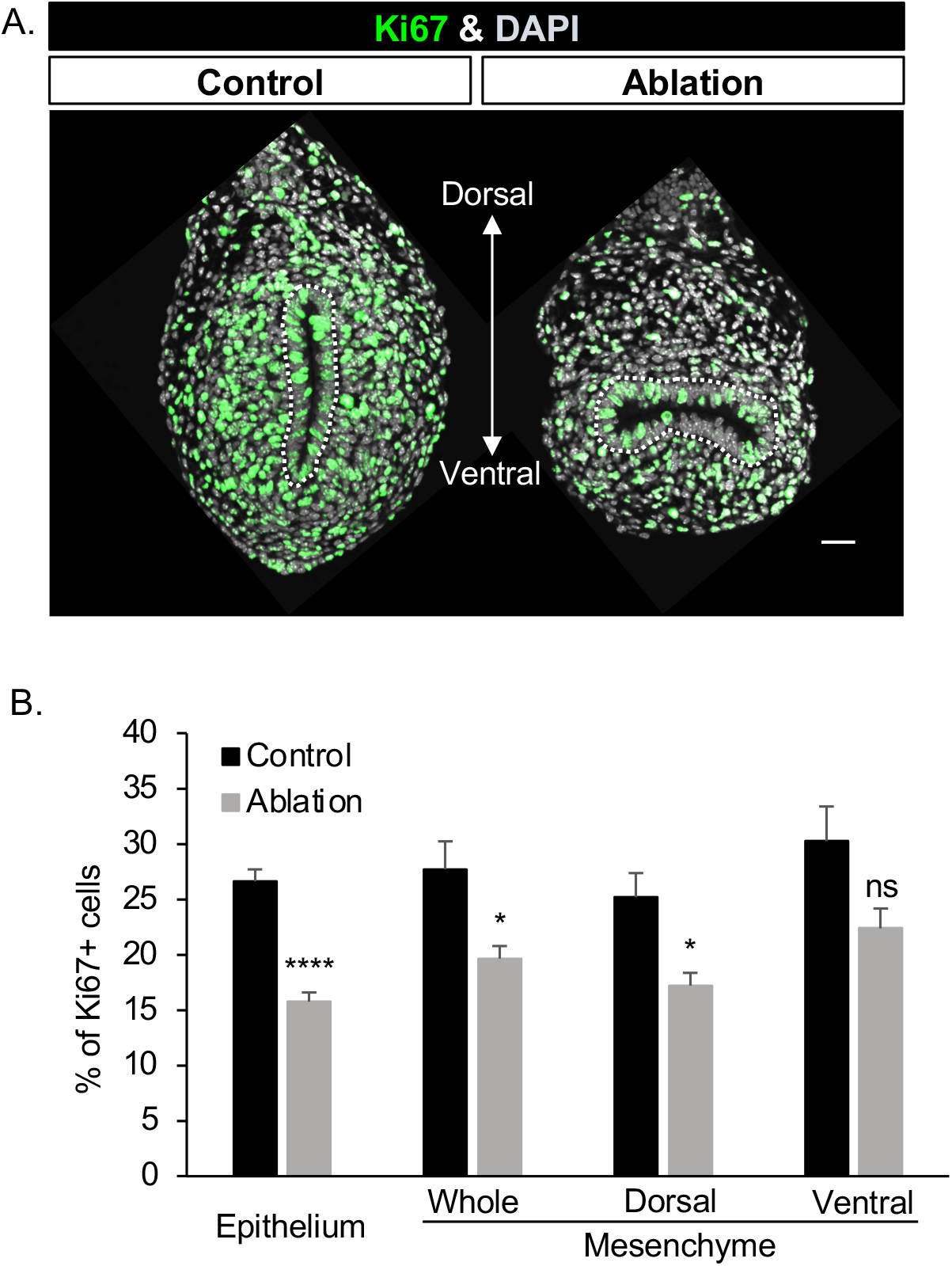
Decreased cell proliferation in epithelial and mesenchymal compartments of the uterus in the ablation group. (A) Representative images of Ki67 immunofluorescence staining on PND0 uteri of the control and ablation groups. Nuclei were stained with DAPI. White dashed circles: uterine lumen. Scale bar: 25 µm. (B) Percentages of Ki67+ cells in the epithelial and mesenchymal compartments of PND0 uteri in the control and ablation groups. N=4-5. Results were shown as mean ± s.e.m. and analyzed by two-tailed unpaired Student’s t test. ns, p>0.05; *, p<0.05; ****, p <0.0001.

### Delayed smooth muscle formation at the dorsal side of the uterus in the ablation group

Smooth muscle formation is a key event of mesenchymal differentiation in the uterus (Gao et al., 2014). To determine whether smooth muscle formation was altered, we performed immunofluorescence of the smooth muscle marker αSMA in the uterus at two timepoints, PND3 when the initial formation of circular smooth muscle layer and PND21 when both circular and longitudinal smooth muscle layers are formed (Gao et al., 2014) (**Fig.7**). On PND3, in the control group, αSMA expression was detected in the circumferentially aligned (i.e. circular) smooth muscle cells of the uterus. (**Fig. 7A**). However, in the ablation group, αSMA expression was absent in the dorsal part of the mesenchymal compartment and only formed a three-quarter circle of the smooth muscle layer (**Fig. 7B**). On PND21, both the inner circular layer and the outer longitudinal layer of smooth muscles were visible in the uterus of both groups (**Fig. 7C & 7D**). These results demonstrate that partial ablation of *Amhr2*+ mesenchymal cells delays the formation of circular smooth muscle in the dorsal region of the uterus.

**Fig 7.**
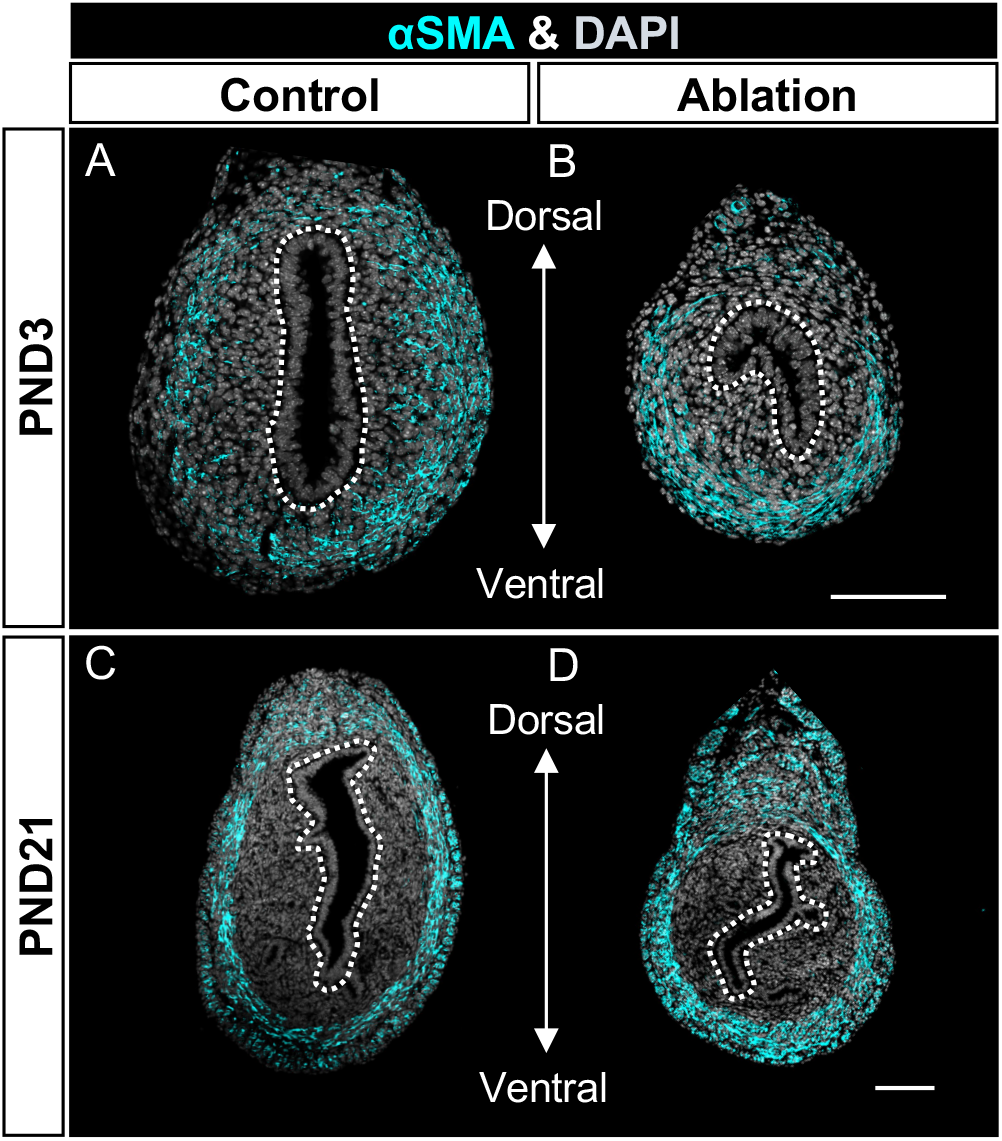
Delayed smooth muscle formation at the dorsal side of the uterus in the ablation group. Representative images of αSMA immunofluorescence staining on PND3 (A & B) and PND21 (C & D) in the control (A & C) and ablation (B & D) groups. White dashed circles: uterine lumen. N=3-4. Scale bar: 100 µm for A-D.

## Discussion

Our work reveals that the proper number of *Amhr2*+ mesenchymal cells is required for normal female reproductive tract patterning along both antero-posterior and dorso-ventral axes. Partial ablation of *Amhr2*+ mesenchyme compromises the oviductal coiling during antero-posterior patterning, alters uterine lumen shape and delays smooth muscle formation in the dorsal-ventral axis.

### Genetic cell ablation represents an important tool for advancing our understanding of female reproductive tract development

Genetic cell ablation technique has been used to discover significant roles of a specific cell type/lineage in development and function of other reproductive organs, including the testis (Caldeira-Brant et al., 2018; Shen et al., 2021; Smith et al., 2015) and ovary (Care et al., 2013; Niu and Spradling, 2020; Turner et al., 2011). However, it has not been used in studying female reproductive tract development and function. Here, we used the *Amhr2-Cre; Rosa-DTA* genetic cell ablation model to reveal consequences of continuous ablation of *Amhr2*+ mesenchyme from the fetal stage on female reproductive tract differentiation. As far as we are concerned, among the phenotypes resulting from partial ablation of *Amhr2*+ mesenchyme, altered uterine lumen patterning and delayed smooth muscle formation have not been reported in any *Amhr2*-Cre driven gene knockouts. Therefore, *Rosa-DTA* model used in our study uncovers new roles of *Amhr2*+ mesenchyme in patterning the epithelial lumen and promoting smooth muscle differentiation. Multiple single cell RNA transcriptomic studies have identified mesenchymal subpopulations and their specific markers in the developing and adult uterus (Kirkwood et al., 2021; Mucenski et al., 2019; Saatcioglu et al., 2019) and in the adult oviduct (Ford et al., 2021; McGlade et al., 2021). With the availabilities of Cre or CreER under the control of marker gene promoters, subpopulation-specific cellular ablation studies can be feasible and potentially provide insights into their specific functions. These observations demonstrate that genetic cell ablation can provide comprehensive roles of a specific cell type/lineage in development and function of the female reproductive tract.

### *Amhr2*+ mesenchyme is essential for oviductal coiling

Tissue recombination studies have demonstrated that the mesenchyme plays inductive roles in epithelial region-specific fates and morphogenesis (Cunha, 1976). However, it is unknown which mesenchymal population directs oviductal epithelial morphogenesis. Using in vivo cellular ablation model, our study demonstrated the critical role of *Amhr2*+ mesenchyme in regulating epithelial proliferation, oviductal elongation and coiling during the antero-posterior patterning. Gene knockout mouse studies have showed that the function of mesenchyme in promoting oviductal coiling is highly dependent upon mesenchyme-derived growth factors. For example, WNT4 is a Wnt ligand that is specifically expressed in Müllerian duct mesenchyme at the fetal stage. Mice with partial loss of WNT4 function presented less-coiled oviducts on E18.5 (Prunskaite-Hyyrylainen et al., 2016).

Interestingly, the distribution pattern of secretory and ciliated cells along the antero-posterior axis of the oviduct was not altered after the partial ablation of *Amhr2*+ mesenchyme. Tissue recombination studies have demonstrated that differentiation of oviductal epithelia into secretory or ciliated epithelial cells is dictated by geographically and functionally distinct oviductal mesenchymes (Yamanouchi et al., 2010). For instance, when recombined with the mesenchyme of the ampulla regions, the epithelia of the isthmus exhibit the distribution pattern of secretory and ciliated cells observed in the ampulla region (Yamanouchi et al., 2010). The normal distribution of the two major epithelial cell types in the oviduct in our cell ablation model suggests that partial ablation of *Amhr2+* mesenchyme does not affect the abilities of region-specific mesenchyme in determining the undifferentiated epithelium to cilial and secretory epithelial type.

### *Amhr2*+ mesenchyme directs uterine lumen patterning along the dorsal-ventral axis

Although *Amhr2+* mesenchyme is a commonly targeted cell population in studying uterine biology, its functions in female reproductive tract development are still not fully understood. Conditional deletion of *Ctnnb1* or *Dicer* in the *Amhr2*+ mesenchyme caused hypoplastic uteri as seen in our *Amhr2-Cre; Rosa-DTA* cellular ablation model *Ctnnb1* (Deutscher and Hung-Chang Yao, 2007; Hernandez Gifford et al., 2009) (Gonzalez and Behringer, 2009). However, these tissue-specific gene knockout mice didn’t show the alteration in the direction of the long axis of uterine lumen. Therefore, our cellular ablation model not only confirms the essential role of *Amhr2*+ mesenchyme for cellular proliferation and tissue growth during Müllerian duct development, but also reveals a novel function of *Amhr2*+ mesenchyme in directing the long axis of uterine lumen. Smooth muscle is known to produce mechanical forces, which is critical for patterning lumen shape (Heisenberg and Bellaiche, 2013; Hinz, 2010). In the ablation group, the delayed smooth muscle differentiation at the dorsal side might reduce mechanical force from the dorsal side for driving/maintaining the direction of the long axis of lumen shape in the dorsal-ventral axis.

AMHR2 is the specific receptor for mediating the action of AMH (Mullen and Behringer, 2014). AMHR2 is expressed in the subluminal mesenchyme from E12.5 through PND6. However, its ligand AMH is not expressed in embryonic and neonatal ovaries until PND6 in mice (Munsterberg and Lovell-Badge, 1991) or PND4-6 in rats(Ueno et al., 1989). Due to the absence of AMH, AMH/AMHR2 signaling remains dormant in the female reproductive tract. Ectopic activation of AMHR2 by exogenous AMH treatment on PND1 via viral gene delivery caused uterine hypoplasia and failure of uterine gland formation (Saatcioglu et al., 2019), which are different phenotypes from those observed in our study (altered direction of the long axis of lumen shape and unaffected uterine gland formation). The phenotypes upon exogenous AMH treatment in their study could result from epithelial death because mesenchymal AMH/AMHR2 signaling is known to induce epithelial degeneration (Mullen and Behringer, 2014). On the other hand, *Amhr2-Cre; Rosa-DTA* cellular ablation model in our study specifically targeted and ablated *Amhr2*+ mesenchymal cells. Therefore, varied phenotypes between these two studies can result from ablating different cell types.

Of note, during embryo implantation, the lumen shape forms a slit-like structure with its long axis in the dorsal-ventral direction, which must be in alignment with the future embryonic axis for establishing successful implantation. Postnatal deletion of *Rbpj* (the nuclear transducer of Notch signaling) in the whole uterus caused tilting of this lumen axis in the Notch pathway-independent manner, resulting in deflected uterine-embryonic axis and substantial embryo loss (Zhang et al., 2014). These results demonstrate the importance of proper establishment of lumen shape orientation and suggest that the uterus with abnormal direction of the long axis in the ablation group could not be able to support successful implantation.

In conclusion, we showed that partial ablation of *Amhr2*+ mesenchyme from the fetal stage impairs female reproductive tract patterning along both antero-posterior and dorsal-ventral axes. The oviduct lost its characteristic coiling due to decreased epithelial proliferation and oviductal elongation. Uterine lumen displayed abnormal shape with its long axis oriented in the left-right direction, perpendicular to the dorsal-ventral axis in the normal condition. Our work opens up the application of genetic cell ablation models for a greater understanding of the function of a specific cell lineage in female reproductive tract development.

## Acknowledgments

We are thankful to Animal Care Staff in the School of Veterinary Medicine for taking care of our mouse colonies.

## Author Contributions

SJ and FZ conceived the study; FZ supervised the project; SJ performed all experiments and analyzed the data. JW and MC analyzed the data. SJ and FZ wrote the manuscript. All authors read and approved the final manuscript.

## Data and materials availability

All data are available in the main text or the supplementary materials upon reasonable request.

---

**Fig S1. Impaired ovarian development in the ablation group.** Representative images of PND21 ovaries in the control and ablation groups N=5 for both control and ablation groups. Scale bar: 1 mm.

**Fig S2. Normal smooth muscle differentiation in the oviduct of the ablation group.** Representative images of αSMA immunofluorescence on the oviducts of the control and ablation groups on PND3 (A) and PND21 (B). N=4, 3 for the control and ablation groups, respectively, in both A & B. Scale bar: 25 µm in A; 50 µm in B.

**Fig S3. Normal uterine gland formation on PND21 in the ablation group.** (A) Immunofluorescence staining of the uterine gland marker FOXA2 (green) in control and ablation groups. White arrows: identified uterine glands. (B) Quantification of uterine gland numbers. N=4 and 3 for the control and ablation groups, respectively. ns: p>0.05 compared to the control group using two-tailed unpaired Student’s t test. Results are shown as mean ± s.e.m. Scale bar: 100 µm.

